# Monitoring mammals, birds and fish during summer 2022 at the outlet of an agricultural stream by mtDNA

**DOI:** 10.1101/2025.05.05.652203

**Authors:** Richard Villemur

## Abstract

Fecal contamination of surface waters poses a potential risk to public and environmental health but can also impact the local economy and recreational activities. Determining its source could facilitate mitigation of the contamination. Fecal contamination can originate from several animals, particularly in areas where urban and agricultural activities overlap. In previous work, we developed a molecular approach to detect the presence of mammals, fish, and birds by sequencing mitochondrial DNA (mtDNA) amplicons derived from environmental DNA. In this report, we monitored the outlet of a stream located in an agricultural area for 16 weeks to detect the presence of mammals, including humans, livestock, domestic and wild mammals, birds and fish. We were able to detect mtDNA sequences affiliated to at least 73 animal lineages. Sequences affiliated to fish were proportionally the most abundant, followed by those affiliated to mammals. We observed increases in bovine and human mtDNA sequences after episodes of high flow in the watershed, suggesting that soil runoff to surface waters carried organic matter (e.g., manure, feces, wastewater) from these animals. Our approach could provide crucial information for farmers to mitigate fecal pollution generated by agricultural activities.

## Introduction

Fecal contaminations are potential risk for public and environmental health but also can have economic consequences on agricultural (irrigation) and recreational activities. Determining its source could facilitate the mitigation of contamination at its point of entry, for example, by restoring faulty infrastructure (e.g., leaking septic tanks, faulty wastewater treatment plants) or improving riparian areas for non-point sources (reducing runoff and improving manure management in the field). However, this determination is challenging because these contaminations are diffuse and can originate far from the sampling locations (McLellan & Eren, 2014, Unno *et al*., 2018). Several source-tracking genetic markers have been developed to determine the animal source of fecal contamination (Harwood *et al*., 2014). For this, DNA extracted from water samples is amplified by polymerase chain reaction (PCR) with primers targeting animal-specific enteric bacteria or animal-specific mitochondrial DNA (mtDNA), which are present in feces. Usually, quantitative PCR (qPCR) tests are performed to quantify the level of these animal host-specific markers, thus assessing the level of water contamination. However, several qPCR assays have to be performed to detect the potential animal(s) responsible for the contamination, which can be cumbersome. Regarding mtDNA as a marker for tracking the source of fecal contamination, detecting the presence of animals from non-fecal matter (skin residues, gray water, feathers or scales) is possible. However, the most abundant animal residues are their excrements (except of course dead animal).

Fecal contamination can be the source of multiple animals, especially in areas where urban and agricultural activities overlap. Furthermore, multiple animal species can be detected in human feces or wastewater (Caldwell *et al*., 2007, Caldwell & Levine, 2009, Kortbaoui *et al*., 2009). For example, the presence of undigested beef in human feces or meat waste flushed down toilets or sinks could be a misleading source of fecal contamination. Although the proportions of these non-human markers in human feces or in human-derived wastewater are much lower than those of the human marker, their presence significantly complicates the interpretation of results when surface waters are vulnerable to fecal contamination by multiple animals. Additionally, wild animals can potentially contribute to fecal contamination, such as wild birds or terrestrial riverbank mammals.

Mitochondrial DNA has become the standard barcoding tool to identify the presence of animal species in different environments. Several PCR primers targeting mtDNA have been developed. So-called “universal” primers with few or no degenerate positions have been developed that target the mitochondrial 12S and 16S ribosomal RNA (rRNA) genes of multiple animals such as mammals and fish (Palumbi *et al*., 2002, Rasmussen *et al*., 2009, Riaz *et al*., 2011, Andersen *et al*., 2012, Giguet-Covex *et al*., 2014, Miya *et al*., 2015). We have developed PCR primers to target the mtDNA of the majority of mammals, birds and fish, and to a certain extent reptiles and amphibians (Ragot & Villemur, 2022). These primers amplify a region of the mitochondrial genome (gene encoding part of the mitochondrial 16S rRNA gene) with a length of about 250 to 300 nt length (amplicon). Using these primers to PCR-amplified mtDNA from environmental DNA (eDNA), they generate an amplicon composed of different sequences representative of the mammals, birds and fish. This amplicon can then be sequenced by by next generation sequencing technologies (e.g. Illumina). In previous study, we used these primers to detect the presence of these animals and determine their proportions in two watersheds (86 sampling sites), in the purpose to assess the potential contribution of fecal contamination of surface waters by multiple animals (Ragot *et al*., 2023).

Using the same approach, we monitored this time one site, the outlet of a stream in an agricultural area, for 16 weeks during summer of 2022 for the presence of mtDNA sequences affiliated mainly to mammals, fish and birds. We discussed the possible contribution of these animals to fecal contamination, as well as the link between the occurring of higher proportions of human and bovine mtDNA sequences following episodes of high watershed flow.

## Material and Methods

### Watershed and water collection

The watershed of the stream is located in the *Baie Missisquoi* region, in a non-urban, agricultural area, around 70 km south-east of Montreal, Canada. It includes several farms. Cattle and pig farms were identified, some with cow grazing. Manure (cattle, pig, poultry) was mainly spread in spring (April, May) and fall (October and November). Two farms spread cattle manure in August (sampling period).

The sampling site was located at the junction of the stream’s outlet to a river. Downstream of the outlet (< 1 km), there is a small marina. The watershed flow expressed as mm/day (m^3^/sec by the watershed area [1237 ha]) was automatically recorded each 15 min. Water sample (1 L) was collected each week, from June 1, 2022 to September 14, 2022 (16 samples), by Mylène Généreux of the *Institut de recherche et de développement en agroenvironnement* (IRDA), and filtered on 0.45 µM filters. Filters were stored at -70°C until use (Table 1).

**Table 1:** Samples and number of sequencing reads.

| Sample Name | Sampling date | Number of reads |
| --- | --- | --- |
| A | 01-Jun-22 | 182 067 |
| B | 07-Jun-22 | 268 566 |
| C | 15-Jun-22 | 252 419 |
| D | 22-Jun-22 | 245 512 |
| E | 29-Jun-22 | 214 649 |
| F | 06-Jul-22 | 259 380 |
| G | 13-Jul-22 | 246 152 |
| H | 20-Jul-22 | 243 340 |
| J | 27-Jul-22 | 310 293 |
| K | 03-Aug-22 | 299 149 |
| M | 10-Aug-22 | 341 864 |
| N | 18-Aug-22 | 309 897 |
| P | 24-Aug-22 | 340 995 |
| Q | 31-Aug-22 | 286 794 |
| R | 08-Sept-22 | 304 684 |
| S | 14-Sept-22 | 342 816 |

Reads are sequences obtained during the sequencing of the amplicons with high quality to make the species affiliations.

### DNA extraction and libraries

DNA extraction from filters and generation of libraries for amplicon sequencing were performed at the Villemur’s laboratory as described (Ragot & Villemur, 2022), with the following modifications in the PCR protocols. The first PCR included 200 ng eDNA (instead of 50 ng) and 0.8 µM (instead of 0.2) of each primer (metaUni126F, metaUni126R; Table 2). The PCR was run at 94 °C for 5 min, then 10 cycles at 94 °C for 30 s, 58 °C for 45 s, 68 °C for 30 s, and 68 °C for 10 min for final elongation. The second PCR included 0.8 µM each of i-metaUni126F and i-metaUni126R (same primers with the Illumina adaptors; Table 2), and 2 µL of the resulted first PCR. The PCR was run under the same conditions of the first PCR with 35 cycles (instead of a touchdown PCR). In the metaUni126F primer described in Ragot and Villemur (2022), nucleotide at third position from the 3’end was changed from a R (A or G) to a D (A or G or T) to generate consensus sequences that include the Otomorpha (Otocephala fish) lineage (Table 2). The resulted PCR products (∼240 nt) was then processed with indexation and cleaning protocols as described in Ragot and Villemur (2022). Amplicon sequencing and sequence analyses are described in Ragot and Villemur (2022). The dada2 pipeline was used to cluster identical reads (ASV). These clusters were then grouped based of 95% identity (named Group ASV), before determining their most probable taxonomic affiliations. Group ASVs that were shorter (∼200 nt) than the expected amplicon length, or with <95% identity, or both, were considered in our analysis. Most of these Group ASVs were related to taxa other than mammals, birds, fish and amphibians, such as insects or crustacean. Raw sequencing data were deposited in Sequence Read Archive (SRA) at the National Center for Biotechnology Information (NCBI: https://www.ncbi.nlm.nih.gov/), Bioproject PRJNA1258561 and Biosamples SAMN48323376 to SAMN48323391.

**Table 2:**
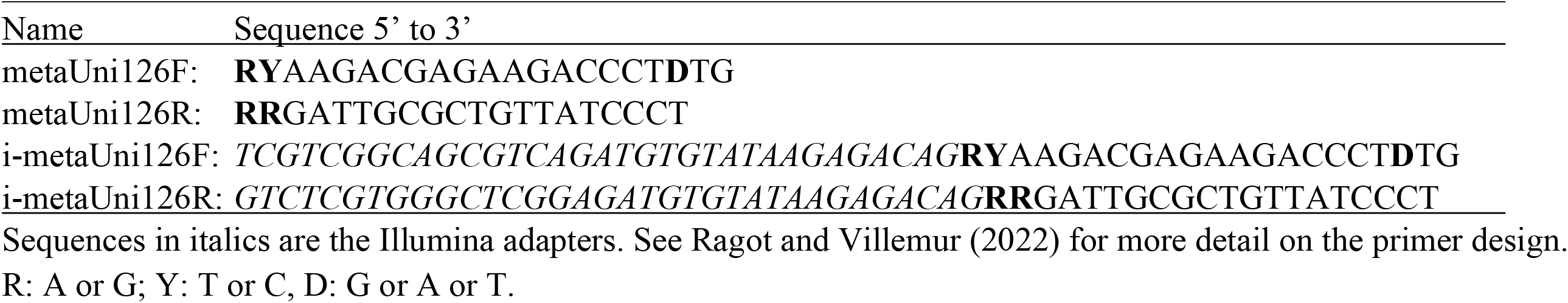
Primer sequences targeting the mitochondrial 16S rRNA gene sequence.

## Results

The total number of reads per sample ranged from 182 067 to 342 816 (Table 1). The most probable affiliations of these sequences were determined with their proportion (Table 3A to F). At least 73 animal lineages were found globally throughout the 16 samples. Figure 1 illustrates the evolution of the relative levels of fish, mammals, birds and amphibians. In average, 61.8% of sequences were affiliated to fish, 29.0% to mammals, 1.0% to birds, 3.6% to amphibians.

**Table 3:** Proportion of animals in water samples. Affiliated reads to specific animal lineages were summed. Results were normalized to *reads by million* to account variability in sequencing depth.

**A-Human, domestic animals and livestock**
|  | June 1 | June 7 | June 15 | June 22 | June 29 | July 6 | July 13 | July 20 | July 27 | Aug 3 | Aug 10 | Aug 18 | Aug 24 | Aug 31 | Sept 8 | Sept 14 |
| --- | --- | --- | --- | --- | --- | --- | --- | --- | --- | --- | --- | --- | --- | --- | --- | --- |
| *Homo sapiens* | 191826 | 269379 | 100767 | 22928 | 69646 | 171166 | 50568 | 170026 | 58864 | 16836 | 85367 | 90474 | 30949 | 156004 | 18493 | 22912 |
| *Bos taurus* | 2233 | 0 | 8300 | 14097 | 195685 | 30093 | 25119 | 27840 | 9293 | 0 | 8521 | 3779 | 133130 | 85138 | 30272 | 4441 |
| *Canis lupus* | 0 | 0 | 14092 | 0 | 0 | 0 | 2373 | 0 | 0 | 1875 | 4781 | 0 | 10902 | 0 | 0 | 935 |
| *Gallus gallus* | 1359 | 0 | 0 | 0 | 0 | 0 | 0 | 60 | 1882 | 0 | 205 | 0 | 12 | 0 | 555 | 121 |
| *Sus scrofa* | 283 | 0 | 915 | 0 | 0 | 0 | 0 | 0 | 0 | 772 | 18 | 255 | 0 | 828 | 0 | 0 |
| *Ovis aries* | 0 | 0 | 0 | 0 | 0 | 0 | 0 | 0 | 0 | 0 | 0 | 0 | 0 | 0 | 0 | 58 |

**B-Wild mammals**
|  | June 1 | June 7 | June 15 | June 22 | June 29 | July 6 | July 13 | July 20 | July 27 | Aug 3 | Aug 10 | Aug 18 | Aug 24 | Aug 31 | Sept 8 | Sept 14 |
| --- | --- | --- | --- | --- | --- | --- | --- | --- | --- | --- | --- | --- | --- | --- | --- | --- |
| *Ondatra zibethicus* | 57998 | 112218 | 126874 | 88923 | 14794 | 83280 | 113621 | 81462 | 213047 | 32103 | 40022 | 102458 | 111942 | 73509 | 215936 | 170659 |
| *Odocoileus virginianus* | 162611 | 14909 | 0 | 35780 | 708 | 28850 | 0 | 15507 | 4066 | 1693 | 44743 | 2377 | 0 | 0 | 1073 | 0 |
| *Microtus* spp. | 568 | 20105 | 0 | 1662 | 9 | 0 | 7802 | 0 | 2092 | 12 | 0 | 0 | 24298 | 15298 | 2946 | 63687 |
| *Procyon lotor* | 1991 | 21872 | 40 | 1776 | 20732 | 4220 | 4836 | 17336 | 0 | 4132 | 3149 | 1704 | 40954 | 4753 | 7939 | 4130 |
| *Condylura cristata* | 6520 | 16597 | 7280 | 2519 | 16371 | 19471 | 0 | 0 | 3435 | 10020 | 12335 | 15720 | 11333 | 5544 | 6236 | 2816 |
| *Blarina brevicauda* | 0 | 0 | 0 | 798 | 0 | 0 | 21694 | 0 | 0 | 0 | 36092 | 0 | 0 | 0 | 125 | 0 |
| *Peromyscus leucopus* | 0 | 0 | 0 | 0 | 0 | 0 | 2021 | 0 | 0 | 0 | 6605 | 891 | 0 | 8748 | 6032 | 0 |
| *Rattus norvegicus* | 0 | 0 | 0 | 0 | 0 | 0 | 0 | 0 | 0 | 0 | 1661 | 0 | 7112 | 8729 | 0 | 0 |
| *Sylvilagus floridanus* | 0 | 9834 | 5015 | 0 | 98 | 0 | 0 | 0 | 0 | 0 | 0 | 0 | 198 | 0 | 0 | 0 |
| Others (5) | 324 | 0 | 42 | 0 | 0 | 0 | 0 | 0 | 58 | 0 | 48 | 0 | 0 | 8783 | 0 | 2698 |

**C-Birds**
|  | June 1 | June 7 | June 15 | June 22 | June 29 | July 6 | July 13 | July 20 | July 27 | Aug 3 | Aug 10 | Aug 18 | Aug 24 | Aug 31 | Sept 8 | Sept 14 |
| --- | --- | --- | --- | --- | --- | --- | --- | --- | --- | --- | --- | --- | --- | --- | --- | --- |
| *Anas platyrhynchos* | 6767 | 28034 | 2266 | 63 | 585 | 8135 | 0 | 857 | 19 | 0 | 2641 | 0 | 7510 | 0 | 143 | 0 |
| *Agelaius phoeniceus* | 1332 | 1819 | 0 | 615 | 1719 | 25056 | 0 | 0 | 3087 | 0 | 0 | 0 | 0 | 0 | 0 | 0 |
| *Turdus migratorius* | 0 | 0 | 0 | 434 | 6438 | 3125 | 3301 | 9158 | 0 | 1254 | 0 | 0 | 7088 | 14 | 0 | 0 |
| *Melospiza melodia* | 0 | 5570 | 0 | 179 | 3722 | 0 | 1008 | 0 | 0 | 690 | 3611 | 0 | 3 | 3065 | 323 | 620 |
| Others (14) | 1810 | 35 | 1329 | 2474 | 648 | 241 | 7227 | 2932 | 572 | 0 | 573 | 1044 | 0 | 471 | 200 | 106 |

D-Fish
|  | June 1 | June 7 | June 15 | June 22 | June 29 | July 6 | July 13 | July 20 | July 27 | Aug 3 | Aug 10 | Aug 18 | Aug 24 | Aug 31 | Sept 8 | Sept 14 |
| --- | --- | --- | --- | --- | --- | --- | --- | --- | --- | --- | --- | --- | --- | --- | --- | --- |
| *Culaea inconstans* | 275207 | 196379 | 259852 | 547060 | 425866 | 437812 | 260179 | 329100 | 315737 | 290103 | 402417 | 278441 | 466911 | 258353 | 512049 | 527825 |
| *Catostomus* spp. | 38272 | 95753 | 22443 | 122589 | 113942 | 98445 | 84119 | 115577 | 189411 | 357023 | 169094 | 211758 | 69432 | 76524 | 82732 | 99227 |
| *Esox lucius/eichertii* | 113 | 15011 | 171812 | 55122 | 77 | 0 | 8958 | 75493 | 107972 | 57794 | 13075 | 31057 | 34 | 23564 | 6528 | 11357 |
| *Semotilus atromaculatus* | 28984 | 20367 | 17723 | 10503 | 2516 | 11497 | 14367 | 17687 | 24177 | 142715 | 10645 | 135084 | 10719 | 21674 | 16087 | 25963 |
| *Luxilus chrysocephalus* | 116164 | 48711 | 22718 | 18394 | 6939 | 16187 | 7963 | 27429 | 475 | 11436 | 5198 | 10332 | 0 | 10347 | 2887 | 10673 |
| *Pimephales promelas* | 14176 | 8629 | 7923 | 16329 | 13503 | 0 | 2123 | 5583 | 11384 | 630 | 487 | 0 | 4477 | 13944 | 41056 | 7643 |
| *Tinca tinca* | 255 | 0 | 0 | 578 | 0 | 0 | 0 | 0 | 6407 | 27328 | 2896 | 65043 | 2977 | 0 | 281 | 9173 |
| *Pimephales notatus* | 741 | 27448 | 62044 | 1106 | 0 | 0 | 7134 | 1631 | 0 | 0 | 8252 | 0 | 0 | 0 | 0 | 370 |
| *Perca flavescens* | 1302 | 9 | 0 | 1806 | 0 | 0 | 26713 | 5731 | 5464 | 25130 | 8302 | 16827 | 0 | 0 | 0 | 5146 |
| *Umbra limi/pygmaea* | 0 | 0 | 0 | 126 | 4601 | 0 | 902 | 32109 | 6110 | 1101 | 1688 | 4032 | 1547 | 23667 | 7482 | 3708 |
| *Ambloplites rupestris* | 0 | 0 | 0 | 0 | 1693 | 0 | 0 | 7855 | 6586 | 7373 | 0 | 11820 | 0 | 8330 | 0 | 3648 |
| *Fundulus diaphanus* | 0 | 14981 | 2478 | 1031 | 6848 | 0 | 0 | 0 | 3732 | 0 | 7953 | 1133 | 0 | 0 | 2947 | 605 |
| *Notemigonus crysoleucas* | 308 | 0 | 0 | 0 | 7505 | 0 | 0 | 0 | 97 | 22 | 0 | 0 | 0 | 22752 | 0 | 79 |
| *Hybognathus regius* | 16197 | 0 | 5778 | 2731 | 7769 | 0 | 0 | 0 | 0 | 0 | 0 | 0 | 0 | 0 | 0 | 0 |
| *Lepomis gibbosus* | 1052 | 592 | 568 | 29 | 1759 | 29 | 288 | 47 | 814 | 247 | 0 | 4974 | 0 | 1145 | 604 | 528 |
| *Cyprinella spiloptera* | 1079 | 0 | 0 | 0 | 0 | 0 | 0 | 0 | 0 | 0 | 0 | 0 | 0 | 0 | 7802 | 1582 |
| *Ameiurus* spp. | 1804 | 289 | 186 | 1155 | 2178 | 0 | 0 | 0 | 127 | 2455 | 228 | 663 | 0 | 0 | 0 | 0 |
| Others (16) | 3188 | 2111 | 222 | 0 | 606 | 216 | 89 | 0 | 151 | 2206 | 224 | 0 | 1548 | 373 | 77 | 0 |

E-Amphibians
|  | June 1 | June 7 | June 15 | June 22 | June 29 | July 6 | July 13 | July 20 | July 27 | Aug 3 | Aug 10 | Aug 18 | Aug 24 | Aug 31 | Sept 8 | Sept 14 |
| --- | --- | --- | --- | --- | --- | --- | --- | --- | --- | --- | --- | --- | --- | --- | --- | --- |
| *Anaxyrus* spp. | 6099 | 38398 | 66568 | 33605 | 37687 | 36294 | 310027 | 29311 | 4017 | 43 | 5473 | 40 | 9398 | 19 | 112 | 772 |
| Others (2) | 124 | 0 | 1248 | 0 | 14 | 253 | 286 | 0 | 0 | 0 | 392 | 0 | 0 | 0 | 0 | 115 |

F-Other lineages
|  | June 1 | June 7 | June 15 | June 22 | June 29 | July 6 | July 13 | July 20 | July 27 | Aug 3 | Aug 10 | Aug 18 | Aug 24 | Aug 31 | Sept 8 | Sept 14 |
| --- | --- | --- | --- | --- | --- | --- | --- | --- | --- | --- | --- | --- | --- | --- | --- | --- |
| *Spizellomycetales* | 16862 | 11824 | 35376 | 6212 | 17205 | 13239 | 27140 | 13602 | 11558 | 2990 | 99240 | 2089 | 40751 | 160669 | 22090 | 16159 |
| *Simulium encisoi* | 8903 | 13189 | 51696 | 0 | 228 | 1723 | 1499 | 974 | 641 | 0 | 0 | 0 | 0 | 0 | 125 | 0 |
| Gastropoda | 13792 | 0 | 436 | 7486 | 15584 | 4750 | 6784 | 11827 | 7087 | 1855 | 2653 | 1662 | 804 | 0 | 184 | 2016 |
| Red algae (*Glaucosphaera*) | 0 | 0 | 0 | 0 | 0 | 81 | 0 | 0 | 0 | 0 | 9676 | 0 | 2023 | 2877 | 0 | 0 |
| Crustacean (*Candona*) | 8953 | 1811 | 0 | 684 | 1645 | 1721 | 1426 | 323 | 114 | 0 | 0 | 0 | 1227 | 1838 | 1118 | 0 |
| *Sciuridae* | 4213 | 0 | 2302 | 0 | 662 | 451 | 0 | 70 | 0 | 0 | 0 | 4934 | 408 | 1524 | 0 | 0 |
| Crustacean (*Cypridopsis*) | 1002 | 3232 | 0 | 365 | 0 | 2770 | 0 | 0 | 329 | 0 | 332 | 410 | 1905 | 307 | 2243 | 131 |
| Uncertain | 5589 | 892 | 1709 | 841 | 19 | 896 | 433 | 473 | 1192 | 160 | 1403 | 1000 | 411 | 1210 | 3325 | 96 |

Recurring reads with weak homology to specific species. Some reads were shorter than the expected amplicon length.

**Figure 1.**
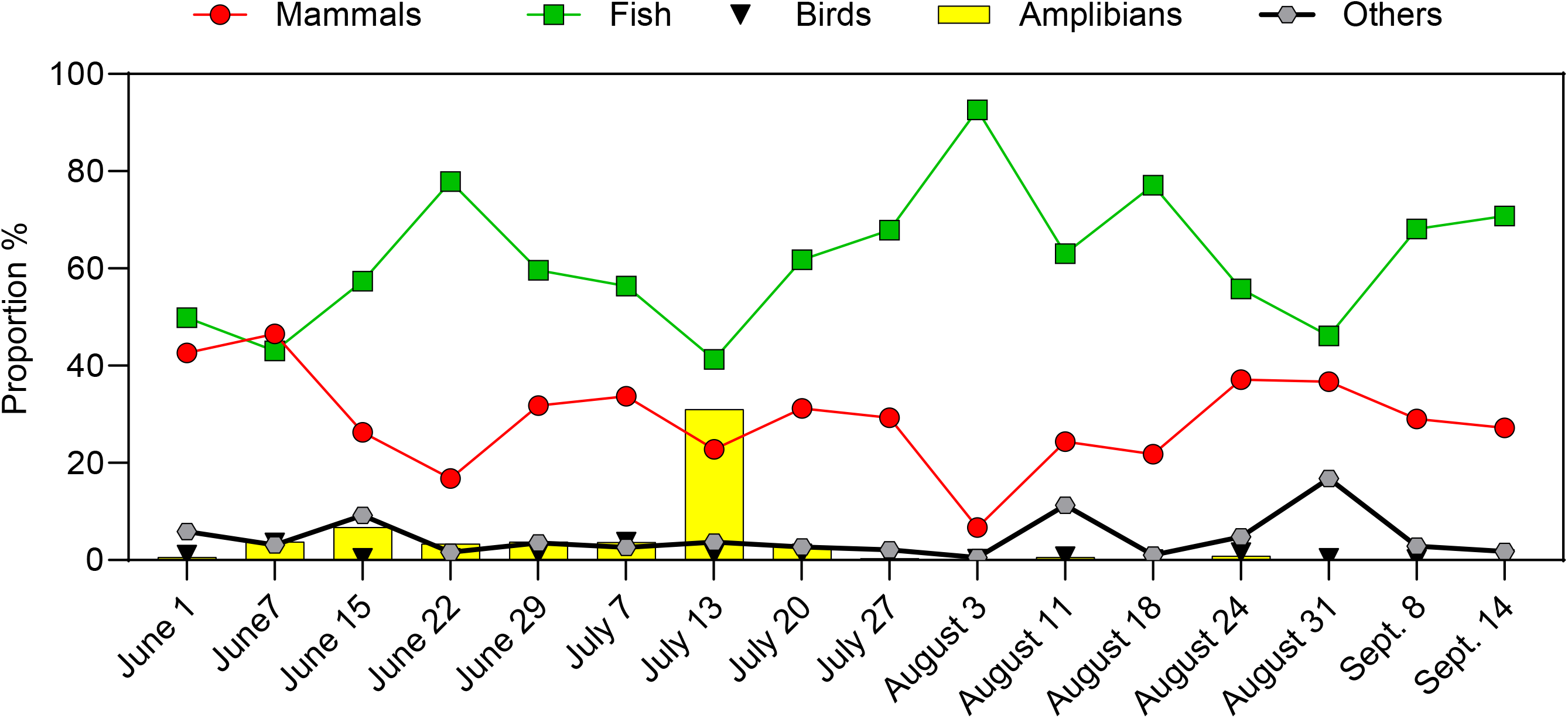
Evolution in the proportions of the mtDNA sequences affililated to different lineages. mtDNA sequences were amplified from eDNA, and amplicon sequencing was carried out. Sequence analysis was performed to affiliate the mtDNA sequences to mammals, birds, fish, amphibians and other lineages.

### Fish

The proportion of sequences affiliated to fish was the highest among the targeted animals, as expected, ranging from 41.3% to 92.6% (Table 3D; Fig. 2). The highest proportion of sequences was affiliated to Brook stickleback (*Culaea inconstans*; 5-cm long fish) ranging from 19.6% to 54.7%, followed by the *Catostomus* spp. (suckers; 20-60 cm long fish) (2.2% to 35.7%).

**Figure 2.**
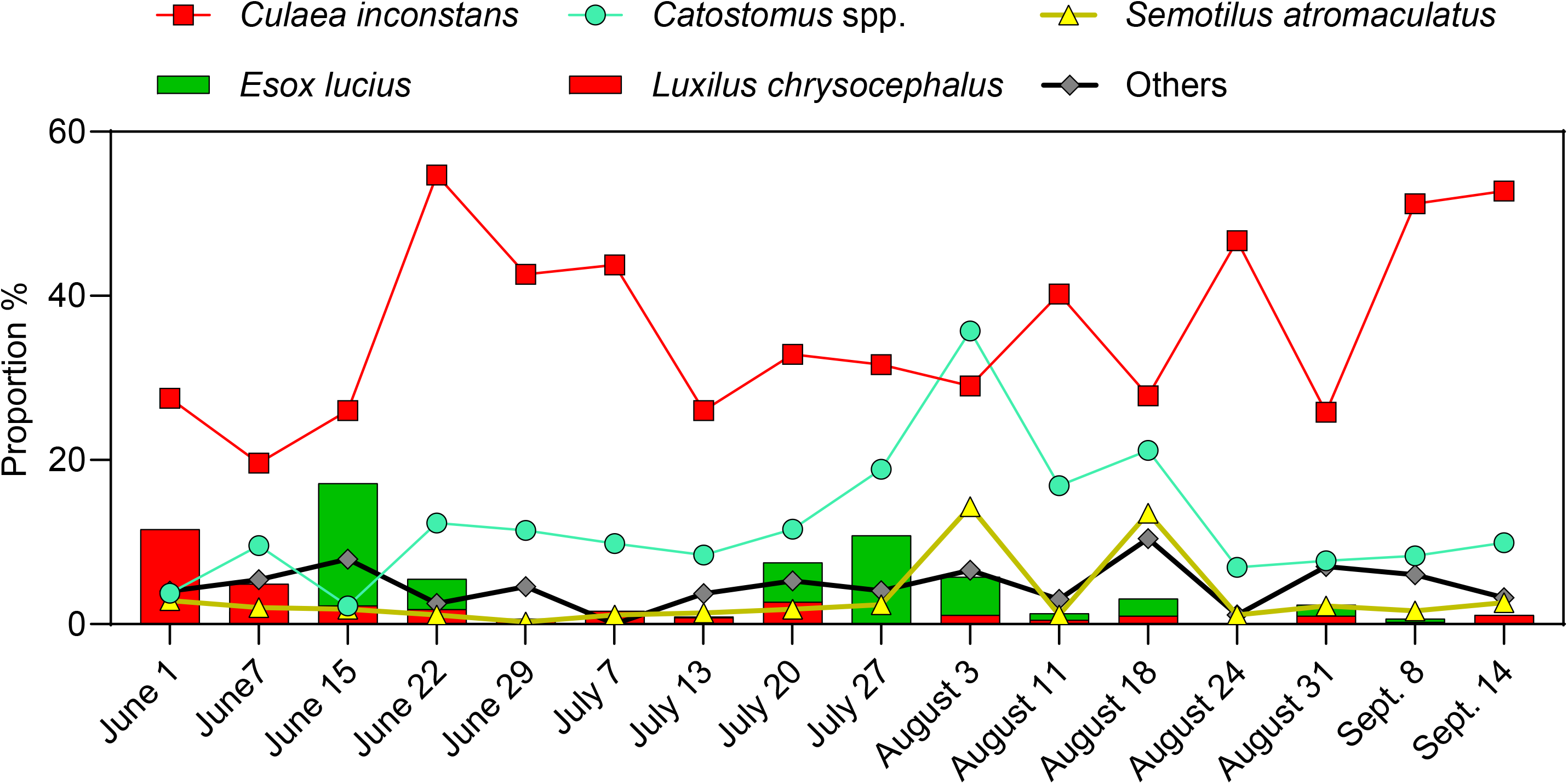
Proportions of the most abundant fish.

### Human, domestic animals and livestock

Human sequences were found in all samples, ranging from 1.7% to 26.9%. Bovine sequences (*Bos taurus*) were detected in 14 out of 16 samples, up to 19.6% (Table 3A). Dog (*Canis lupus*), poultry (*Gallus gallus*), ovine (*Ovies aries*) and porcine (*Sus scrofa*) sequences were scarcely detected in low proportions (Table 3A).

### *Wild mammals and birds* (Table 3B and C)

Muskrat (*Ondatra zibethicus*) sequences were found in all samples, ranging from 1.5% to 21.3%. White-tailed deer (*Odocoileus virginianus*) sequences were found in 11 out 16 samples, one at 16.3% on June 1 (first sample). Racoon (*Procyon lotor*) sequences were also found in most samples at the highest proportion (4.1%) in the August-24 sample. Although sequences affiliated to several bird taxa (18) were detected, they were found in low proportions compared to mammals and fish, ranging from 0.07% to 3.7%.

### Other taxa

Amphibians (Table 3E), especially toad and frog (*Anaxyrus* spp.), were found in higher proportions (up to 31.0%) in first part of summer (June to mid-July). We noticed a Group ASV with shorter sequences affiliated to the fungi *Spizellomyces* (Table 3F). It was first ruled out of the analysis as chimera, but because of the high proportions of these sequences (found in all samples up to 16.1%), they were kept as potential organisms detected by our approach. Although PCR conditions were relatively stringent (58°C hybridization temperature), possible sub-optimal hybridization of our primers to mtDNA of fungi, insects and non-target animals has probably happened. For instance, the highest proportion of sequences affiliated to black fly (*Simulium encisoi*) occurred early in June, then were almost completely absent afterwards (Table 3F), which concurs with the life cycle of these annoying bugs. However, due to sub-optimal conditions of the mtDNA amplification of those organisms, their relative levels may not be accurate.

### Watershed flow

Figure 3 illustrates the evolution of the most prominent mammals (human, bovine, muskrat and white-tail deer) during summer as a function of the watershed flow. Three episodes of high flow observed in late May, June and August were representative of recent rainfall events that occurred prior these episodes and could have generated runoff from soil to surface waters. Higher proportions of human sequences were observed after each of these episodes. However, higher proportions of human sequences were also observed (mid-July, mid-August), unrelated to high flow episodes. Proportions of bovine sequences increased substantially after the second and the third high flow episodes. In addition, higher proportion of bovine sequences was also observed in mid-August prior to the high flow episode, which may have been caused by the spreading of cattle manure by two farms at that time. Fluctuations in the proportions of the muskrat sequences did not show any relationship with the watershed flow.

**Figure 3.**
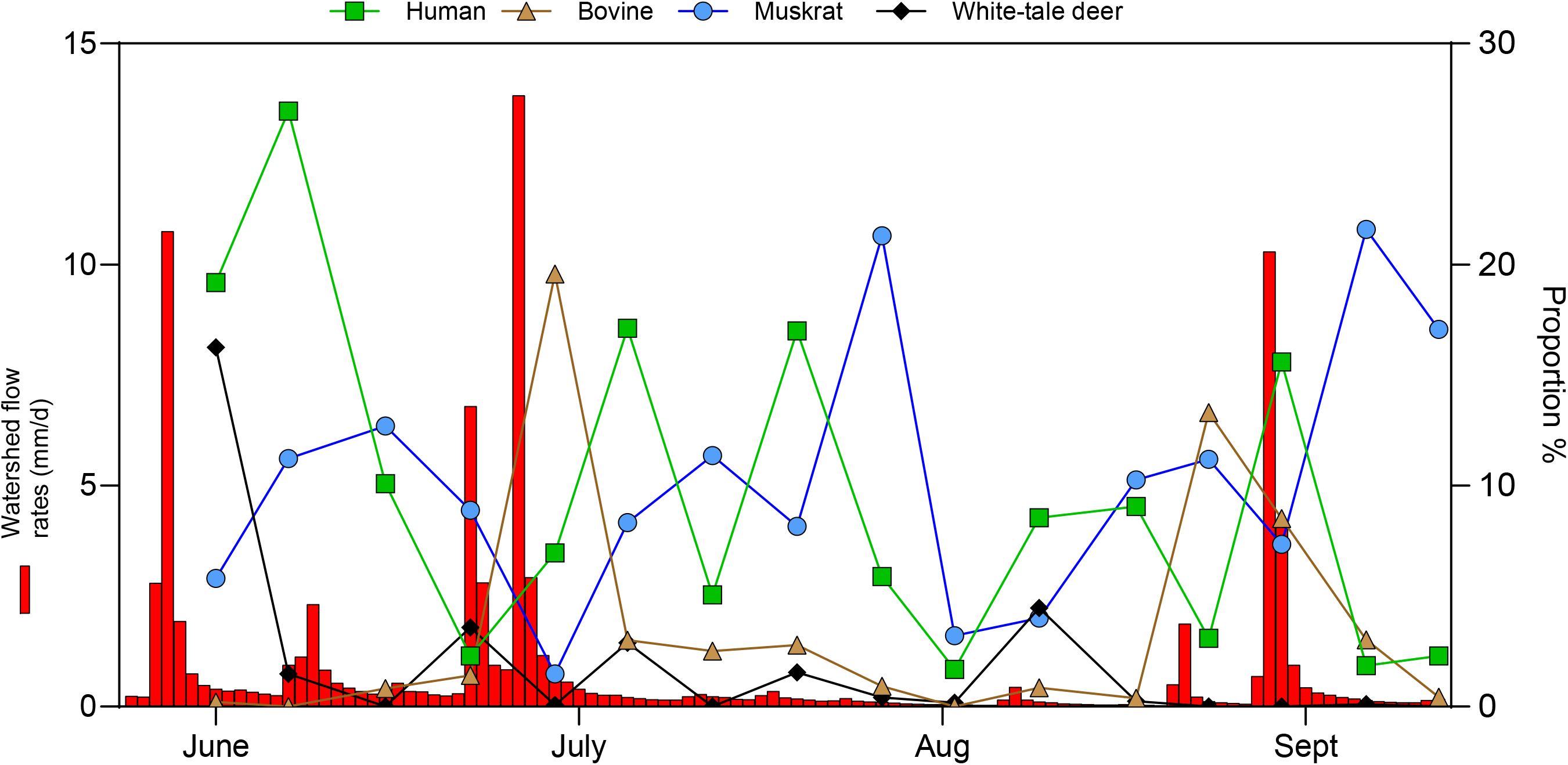
Proportions of the most abundant mammals in the samples, in relation to the watershed flow. The proportions of grouped asvs affiliated to human, bovine, muskrat and white-tale dear are illustrated. Watershed flow was recorded each 15 min. Each bar represents the average of 100 consecutive measurements.

## Discussion

Surface waters contain materials composed of (among other things) suspended organic elements. These organic elements come from the biomass of various sources, including living or dead organisms (algae, plants, microorganisms), or from the waste of larger animals living in the water (fish, amphibians) or in the surrounding area (mammals, birds, reptiles). These wastes can be of different natures: excrement, skin, saliva, urine, feathers, dead animals in decomposition, sperm and fish eggs during spawning. In addition, improper connection of sanitary sewers, defective septic tanks or overflow events can lead to the presence of human and pet feces, as well as greywater and meat waste consumed by the population (pork, beef, poultry, fish).

The organic elements trapped on the filters were extracted to purify total DNA (eDNA). This DNA represents a snapshot of the diversity of organisms present, or that were present, in the water or surrounding areas at the time of sampling. Free DNA trapped in the sediment could also have contributed to the extracted eDNA. For example, animal feces or manure spread on the field could have been washed into surface waters by a rainstorm. This eDNA contains, among other things, the genomes of these organisms in addition to their mtDNA.

Our approach provided a good estimate of the mammals, birds, and fish present over the 16 weeks, and their proportion. The higher proportions of bovine sequences in June and August are correlated with episodes of high watershed flow that followed significant rainfall a few days earlier (recorded by Environment Canada stations). Although there was no manure application in June, cattle grazing was present, which could explain the higher proportion of bovine sequences. In August, in addition of cattle grazing, two farms applied cattle manure, which is also consistent with higher proportions of bovine sequences in mid-August. Other studies reported higher levels of fecal indicator bacteria after after rainfall or high flow events (McDaniel *et al*., 2013, Yakirevich *et al*., 2013, Liao *et al*., 2015, Balkhair, 2017), which concur with our results with mtDNA.

Higher proportions of human sequences occurred after the three episodes of high watershed flow (May, June and August). However, these relative levels also fluctuated when watershed flow was low. Faulty wastewater treatment plants (e.g. septic tanks) could have caused recurrent discharge of human matter into surface waters. In addition, presence of a small marina nearby, and swimming activities could have contributed to these human discharges in surface waters.

In a recent work, Saleem et al. (2025) used a similar molecular approach in monitoring mtDNA in urban areas. They detected mtDNA sequences affiliated to several mammals and birds, including human, livestock and wild animals. By combining these data with microbial source tracking markers, the authors provided a more comprehensive characterization of potential sources of fecal contamination.

It is important to understand that the quantitative aspect reveals proportions, not absolute concentrations, due to the nature of sequencing. Thus, human-related sequences may be highly abundant compared to other species in one sample, but very low in another sample. However, in absolute terms, the number of human-related sequences could be identical in both samples. It is the other species in the first sample that are less abundant in absolute terms, and much more abundant in the second sample. Only qPCR assays can estimate these absolute values.

Another aspect of the quantitative data must be taken with caution. A high proportion of one animal relative to another may be related to the size of these animals. Thus, cows (for example) are certainly less numerous than small fish present in the watershed. It is just that cow excrements are more abundant per individual than those of a small fish.

An interpretation of the results that should also be considered with caution is whether the individual animal excrements were recent (a few days) or recurrent (daily). The excrements may have occurred several days or even weeks earlier and underwent degradation over several days or were trapped in the sediment. In this case, the level of mtDNA of these excrements does not allow to estimate their importance at the time of the excretion. It can be assumed that recurrent droppings from animals endogenous to the water body, including fish, amphibians, or aquatic mammals, are expected. The distance between the animal droppings and the sampling site must also be taken into account. Thus, excrements from cows (for example) a few kilometers upstream from the sampling site will be less abundant on the site (due to dilution and degradation) than those from cows present in the surrounding area of this site. It must also be taken into account that some animals are migratory, i.e., they are present at one time of the year and absent the rest of the time (e.g., birds). In addition, the life cycle of an organism may, at a certain time of year, release more excrement (fish and spawning grounds). Sampling the site for 16 weeks allowed us to better identify animals that permanently inhabit the watershed, rather than those that are transient or those that develop during this period. For example, sequences affiliated with the Brook stickleback were present in high proportions throughout the summer, except in August with a high proportion of *Catostomus* spp. (suckers), which suggests an increase in the development of this fish during this period.

## Conclusions

We were able to assess the presence and the proportion of mammals, birds, and fish (and much more) over a 16-week period at the outlet of a stream in an agricultural area. This information is of great interest to farmers, as it provides insights into the impact of agricultural activities (grazing, manure spreading) on surface water quality. Our results also established a correlation between high flow episodes in the watershed and higher proportions of bovine and human mtDNA sequences. Globally, regular monitoring of watersheds using our approach could lead to better agricultural practices and better maintenance of wastewater treatment infrastructure.

